# NeuroFold: A Multimodal Approach to Generating Novel Protein Variants *in silico*

**DOI:** 10.1101/2024.03.12.584504

**Authors:** Keaun Amani, Michael Fish, Matthew D. Smith, Christian Danve M. Castroverde

## Abstract

The generation of high-performance enzyme variants with desired physicochemical and functional properties presents a formidable challenge in the field of protein engineering. Existing *in silico* design methods are limited by inadequate training data, insufficient diversity within datasets, and suboptimal sampling techniques. Here, we introduce a novel approach that addresses these limitations and significantly improves the efficiency of generating functional enzyme variants. Using a multimodal approach, NeuroFold can leverage sequence, structural, and homology data during both sampling and discrimination phases, thereby enabling more diverse and informed sampling of the sequence space. Our model demonstrated a 40-fold increase in Spearman rank correlation as compared to large language models (LLMs) such as ESM-1v and empowers the rapid creation of high-quality enzyme variants, such as the β-lactamase variants generated by NeuroFold in this study, which demonstrated increased thermostability and varying levels of activity. This pipeline represents a promising advancement in the field of enzyme engineering, offering a valuable tool for the development of novel enzymes with enhanced performance and desired chemical properties.

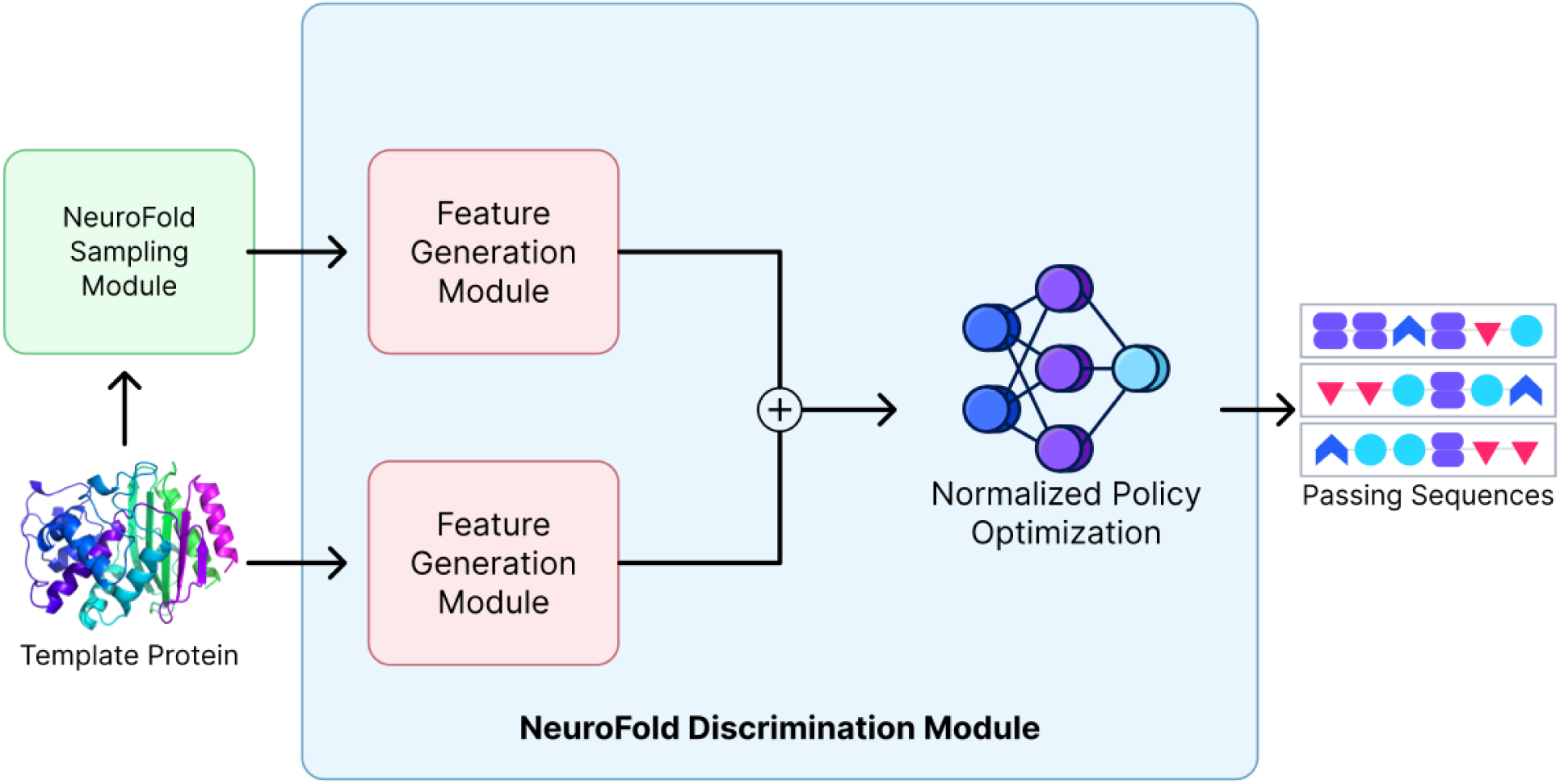

## Introduction

Enzymes, as proteins that facilitate biochemical reactions, possess remarkable practical and industrial significance (Robinson, 2015). The ability to create customized enzyme variants with specific chemical properties, such as enhanced thermostability, increased or decreased reaction rate, broader pH stability, and improved solubility holds immense value in various applications (Li et al., 2020). However, generating such desired variants remains an exceptionally difficult task, primarily due to the magnitude of the protein sequence space (Romero & Arnold, 2009).

Conventional experimental methods, like deep mutational scans (DMS), prove undesirable for generating improved enzyme variants as they necessitate costly and highly parallelizable assays to be effective while only sampling a small portion of the protein sequence space, typically limited to a few point mutations or indels (Notin et al., 2022). For example, the fitness landscape of point mutations in three indole-3-glycerol phosphate synthase (IGPS) orthologs was assayed in one study to reveal the importance of both sequence and structural features of these model TIM barrels (Chan et al., 2017). Another study used multiplex assays to assess the differential abundance and activity of missense variants of the Vitamin K epoxide reductase (VKOR) enzyme (Chiasson et al., 2020). As demonstrated by these and other studies, the production of a substantial number of high-quality enzymes is severely restricted by the above-mentioned limitations, such as time-/labor-intensive assays and/or sub-universal protein sequence coverage.

Recently, the emergence of deep learning-based models has led to the development of novel *in silico* techniques for designing enzyme variants. Protein language models (pLMs) such as ESM-MSA (Rao et al., 2021), ESM-1v (Meier et al., 2021), and CARP-640m (Yang et al., 2022) have demonstrated varying levels of success in enzyme fitness prediction. These newer approaches offer the ability to generate better enzyme candidates more rapidly while streamlining the experimental bottleneck required in the past. Despite their advantages, deep learning models also have their limitations, including inconsistent accuracy and an inability to generalize effectively to certain proteins such as large and/or multimeric enzymes (Table 1).

**Table 1.**
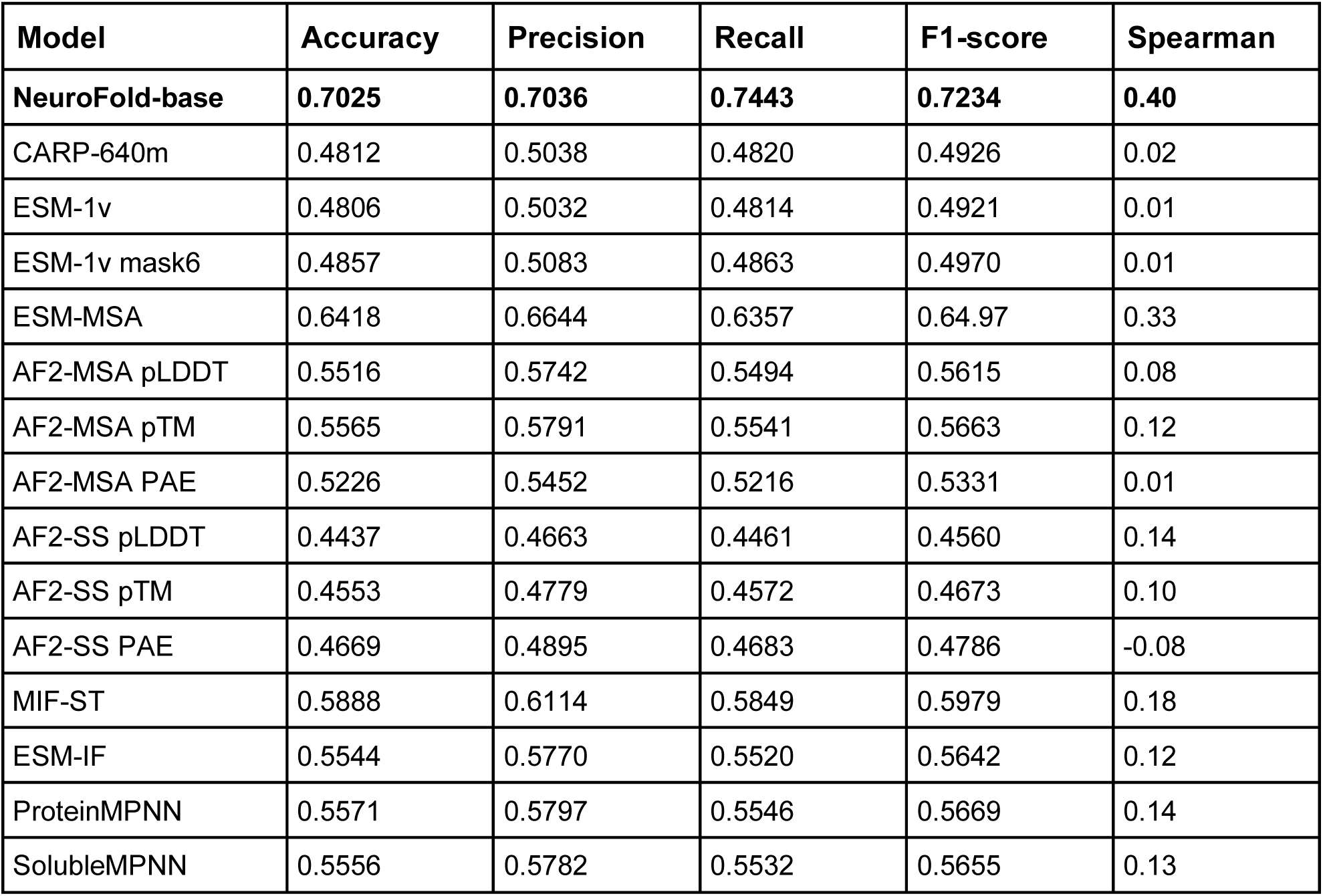
Evaluation of NeuroFold’s performance using our dataset and comparison to various other models. Prediction accuracy, precision, recall, F1-score, and Spearman rank correlation with activity / fitness were calculated. The cutoffs used for all models was the sample 50th percentile. AF2-MSA corresponds to ColabFold with mmseqs2 MSA generation while AF2-SS corresponds to ColabFold with single sequence input.

Structural models such as AlphaFold2 are still unable to consistently predict mutations that impact protein structure (Table 1). Additionally, while single sequence methods like ESM-1v and CARP-640m have demonstrated reasonable accuracy for discriminating against mutations, they can sometimes struggle with orphaned sequences (proteins with low or no homology to known domains or proteins) (Table 1). We hypothesize that the limitations of ESM-1v, CARP-640m and related models are due to single sequence models being unable to capture co-evolutionary relationships that are not implicitly distilled into their model weights. This concept is reinforced further by evolutionary methods such as ESM-MSA, which demonstrate significantly greater performance when it comes to discriminating against mutations (Table 1). Arguably, this is due to being explicitly given an input multiple sequence alignment (MSA) from which to extract co-evolutionary relationships, as well as being able to implicitly capture information related to the protein landscape within their model weights.

In the current study, we present NeuroFold, an innovative hybrid model for designing enzyme variants with desired traits that incorporates a unique architecture to address the limitations of previous methods. Unlike conventional approaches that predominantly rely on a single modality such as sequence or evolutionary data in the form of an MSA (Rao et al., 2021), NeuroFold goes beyond reliance on a single modality by integrating sequence and evolutionary data with structural information. By combining these diverse data sources into a single multimodal approach, our model is able to better reason about the protein space, enabling the inclusion of theoretical constraints that guide the generation of viable enzymes. As a proof of principle, Neurofold was successfully used to design variants of β-lactamase with enhanced thermostability and varying levels of activity relative to the wild-type. As a result, NeuroFold achieves a broader coverage of the sequence space and demonstrates the potential for confidently producing enzyme variants with improved stability, desirable physicochemical properties, and enhanced reaction rates.

## NeuroFold Architecture

### Sampling Module

The NeuroFold architecture starts with an input amino acid sequence that acts as the template or reference. This template sequence should correspond to a functional enzyme that can also be used as a baseline for comparison (e.g., the wildtype version of an enzyme). This sequence can then be used to produce a large number of variants using either the MSA approach, an inverse folding approach, or a combination of the two methods (Figure 1).

**Figure 1.**
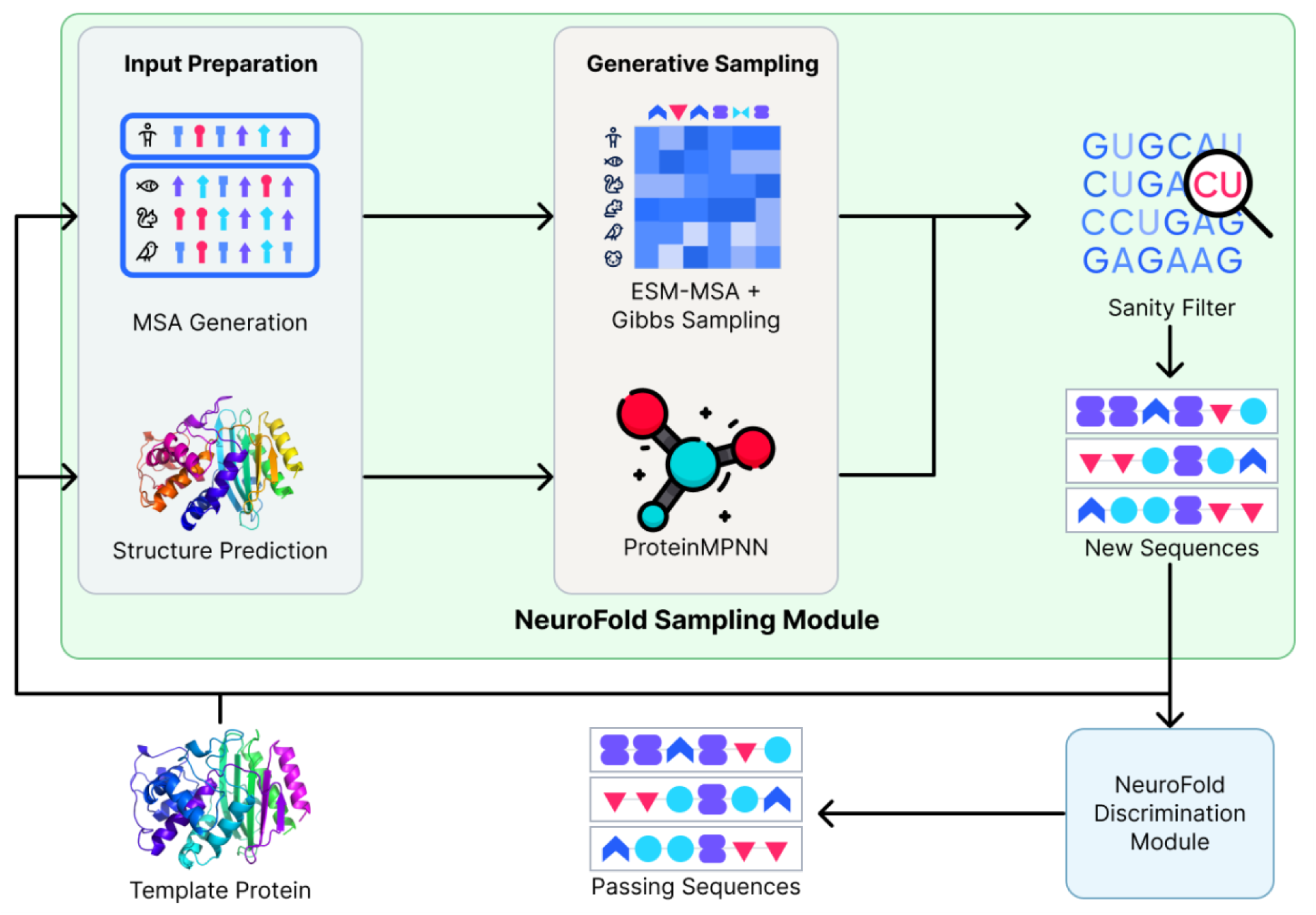
Outline of NeuroFold’s Sampling Module. NeuroFold’s architecture can be broken down into 3 core modules: The Sampling Module, Discrimination Module, and Feature Generation Module. The Sampling Module is responsible for producing the candidate sequences. The Discrimination Module is responsible for discriminating against functional and non-functional candidates generated by the sampling module. Finally, the Feature Generation Module is a component of the Discrimination Module that is responsible for producing the input features of each candidate sequence.

The MSA approach consists of using ESM-MSA (Rao et al., 2021) with Gibbs sampling (Johnson et al., 2021) to produce novel sequences from an input MSA produced from the template sequence. This method is useful for sequences with rich homology and allows novel sequences to be inferred from evolutionary data (Rao et al., 2021).

The alternative inverse folding approach utilizes ProteinMPNN (Dauparas et al., 2022) to generate new sequences from a template structure. This is ideal when a high quality experimental structure is available for the template protein and was employed in this case (see Table S1 for settings). Inverse folding operates on the assumption that proteins with similar structures will tend to possess similar properties, an assumption that is supported by our findings.

The benefit of utilizing an inverse folding approach is that models such as ProteinMPNN are typically capable of sampling a much broader sequence space with sampled sequences, typically sharing 40-70% pairwise sequence identity to the wildtype. This can be a powerful advantage over MSA and traditional approaches such as DMS, as proteins with divergent sequences can still produce similar folds and even carry out similar functions. For example, Koehl and Levitt (2002) showed that the B1 domain of the streptococcal Protein G and *P. magnus* Protein L share striking structural similarities in spite of their low amino acid sequence identity (Figure S1).

The sequences generated from the sampling models are then evaluated using a static sanity filter consisting of rules such as removing sequences that contain an excessive number of repeats as well as removing sequences that contain disruptive or rigid amino acids in loop regions. The new set containing the filtered sequences is then passed into the Discrimination Module alongside the original template sequence. The result is a subset of sequences predicted to be fit by the Discrimination Module. These sequences can then be experimentally validated or used for additional downstream evaluation.

### Discrimination Module

The Discrimination Module (Figure 2) is responsible for determining whether or not a protein variant is fit with relation to a reference protein. This module receives two input sequences, a template sequence that acts as the reference and a mutated or variant sequence that acts as the candidate to evaluate. The template sequence is critical as it grounds the model with a baseline to compare as a reference. By introducing this baseline, the model learns to compare and measure the difference in potential fitness compared to just learning an arbitrary relative fitness for the entire protein landscape.

**Figure 2.**
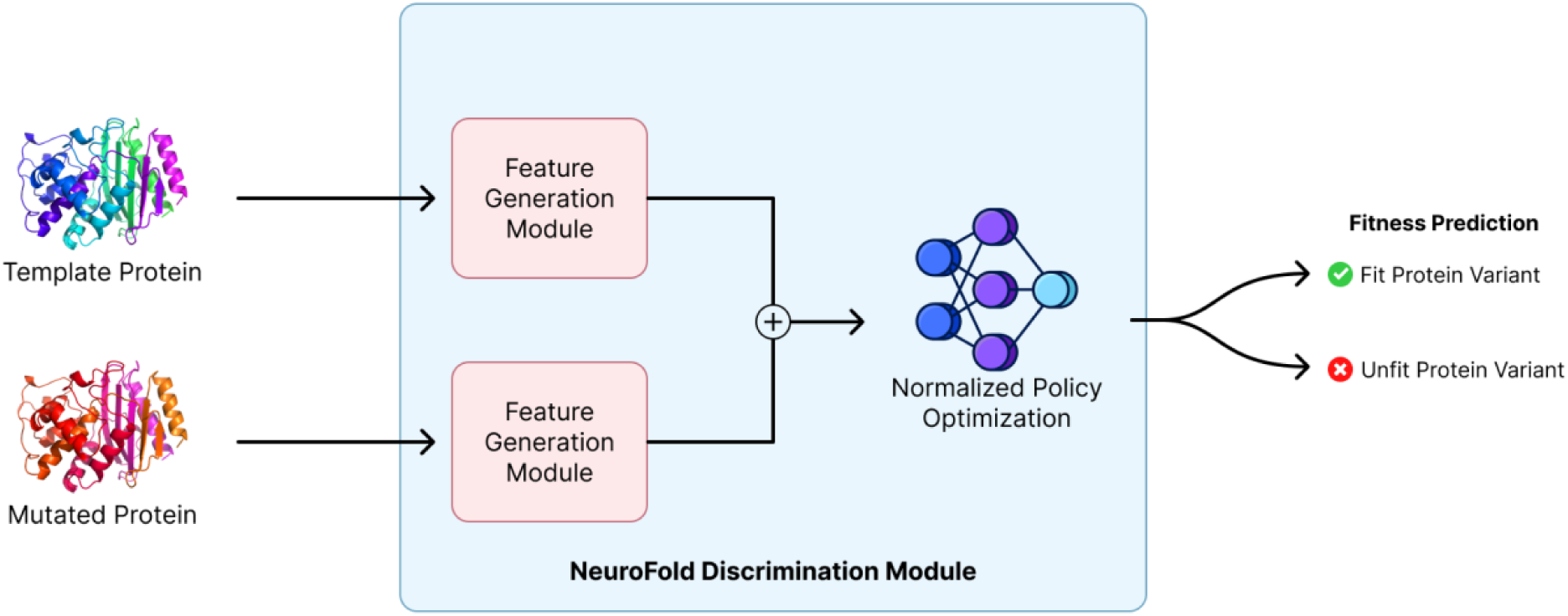
Outline of NeuroFold’s Discrimination Module. The Discrimination Module is responsible for differentiating between a fit and unfit enzyme. The module itself effectively acts as a binary classifier that predicts whether an enzyme can be considered fit.

These sequences are then passed into the Feature Generation Module (Figure 3) to produce two sets of corresponding output representations. These output representations are then fed into a Normalized Policy Network (NPN) which classifies the mutated sequence as either fit or unfit relative to the template sequence. The NPN acts as a binary classifier using the output representation from the FGM as input.

**Figure 3.**
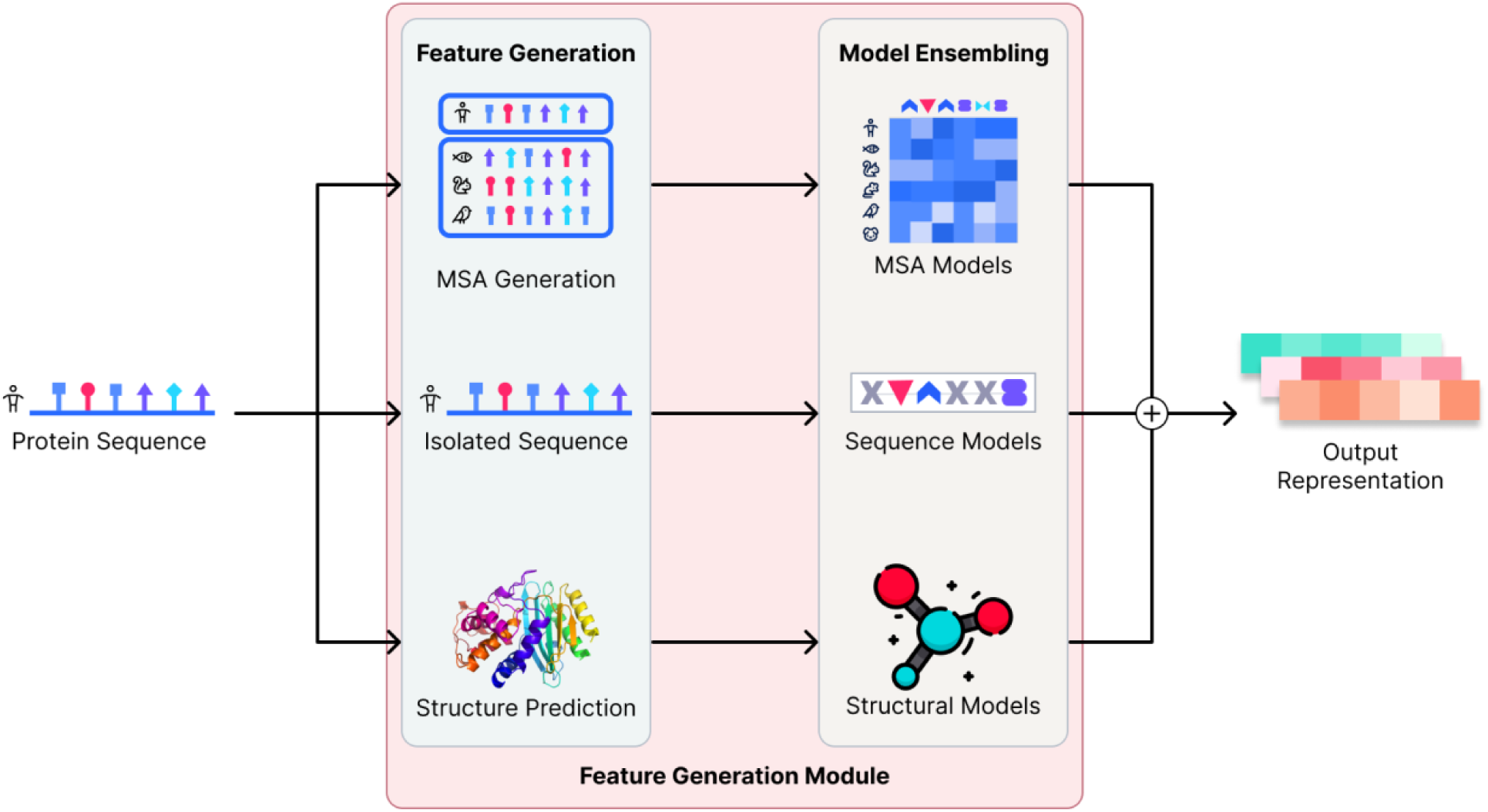
Diagrammatic representation of the NeuroFold Feature Generation Module (FGM). The FGM uses a multimodal approach to capture meaningful representations from sequence, evolutionary, and structural data.

### Feature Generation Module

The Feature Generation Module (FGM) is a key part of what gives NeuroFold its predictive power. The FGM receives an input amino acid sequence and generates a corresponding output representation which is used throughout the rest of the network. The FGM itself can be split into two main segments. The first segment is responsible for producing features from the input sequence in the form of an MSA and a predicted structure. These features are then passed into the Ensembling Module, a collection of models for each modality. The output from the models are then ensembled by modality and returned as an output representation (Figure 3).

The MSAs are generated locally using the MMseqs2 algorithm (Steinegger, 2017) with UniRef30_2022 as the database (Süzek et al., 2007; Mirdita et al., 2022). For each input sequence, two protein structures are predicted using AlphaFold2-ptm (Jumper et al., 2021; Mirdita et al., 2022); the first structure is predicted using the same generated MSA, while the other is predicted as a single sequence and without MSA as AlphaFold2 and RoseTTAFold (Baek et al., 2021) have both demonstrated higher accuracy on certain sequences when no MSA is provided (Baek & Baker, 2022). This enhanced representation biases the model to produce more realistic proteins that other models without the same constraints might ignore.

### Performance Metrics

NeuroFold was tested on experimentally validated proteins from the ProteinGym dataset (Notin et al., 2022), TAPE stability dataset (Rao, 2019; Rocklin et al., 2017), as well as additional datasets collected from various studies (Johnson et al., 2023; Madani et al., 2023; Repecka et al., 2021; Russ et al., 2020). The subset of proteins used from the ProteinGym dataset was carefully curated so as to exclude experimental data from proteins with irrelevant non-enzymatic functions in processes such as viral replication and protein-protein interactions.

The final dataset consisted of 116,321 individual protein sequences from 38 protein families/taxa (Johnson et al., 2023; Madani et al., 2023; Notin et al., 2022; Rao, 2019; Repecka et al., 2021; Rocklin et al., 2017; Russ et al., 2020), including variants with single point mutations, multiple point mutations, and indels. The benchmarks for fitness within these datasets consist of enzyme activity, enzyme stability, and fluorescence. In addition to NeuroFold testing, the collated experimental datasets were also fed into currently existing models, such as CARP-640m (Yang et al., 2022), ESM-1v (Meier et al., 2021), ESM-MSA (Rao et al., 2021), AlphaFold2 (Jumper et al., 2021), MIF-ST (Yang et al., 2022), ESM-IF (Hsu et al. 2022), ProteinMPNN (Dauparas et al., 2022), and SolubleMPNN.

NeuroFold outperformed all other models with the highest accuracy, f1-score and precision metrics, all of which were higher than alternative approaches (Table 1). For practical applications within the task of enzyme stability prediction, precision is the most valuable metric as it is not the number of false negatives that are of value, but the number of false positives. Since the protein space is so vast, the major limitation is rarely computationally sampling viable sequences. Instead, the bottleneck is typically the experimentation and validation of proteins. Minimizing false positives effectively minimizes the experimental time, cost, and resources.

NeuroFold demonstrated an accuracy of 70.25% when validated on our dataset (Table 1). The next best-performing model was ESM-MSA with 64.18%, with ESM-1v and single sequence AlphaFold2 scoring the lowest in this metric (48.06% and 44.37% respectively). Overall, the accuracy achieved by NeuroFold represents a significant improvement over the typical success rates of ∼14.17% achieved using traditional methods (Notin et al., 2022; Table S2). The precision of NeuroFold on each dataset was also independently computed (Figure 4). We find the overall performance of AF2-SS to be significantly lower than that of AF2-MSA and arguably redundant. For this reason we would recommend excluding this method in the future.

**Figure 4.**
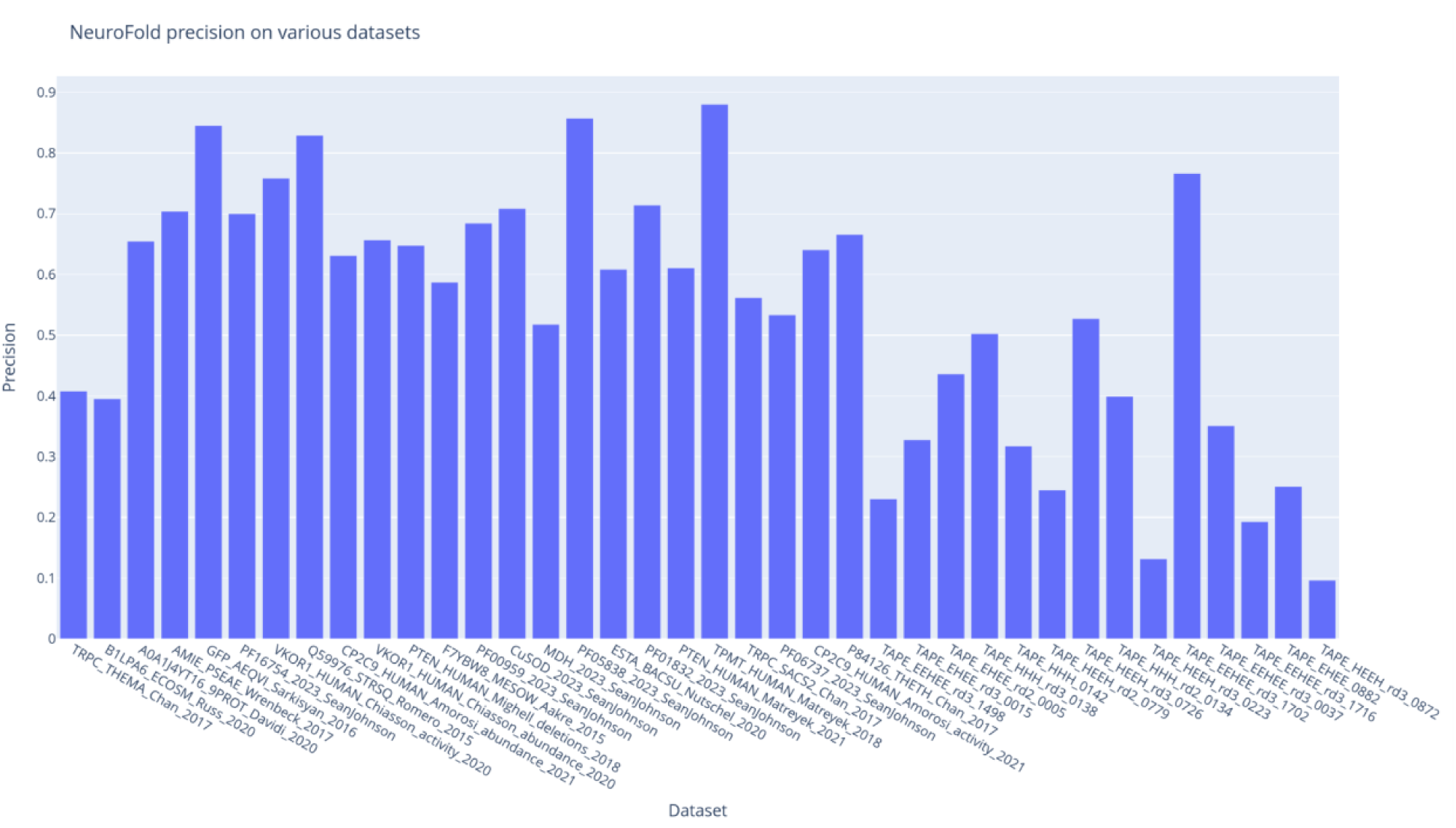
NeuroFold precision on various datasets. NeuroFold’s precision was calculated on 38 different datasets of experimentally validated proteins. A high precision score indicates that NeuroFold correctly predicted the fitness of proteins tested within a specific dataset, while a low precision score indicates a smaller degree of correct predictions per protein variant. The details, computed metrics, computed MSAs, as well as all computed structures are all freely available to download from the following URL https://neurosnap.ai/neurofold/dataset.

#### Model performance on benchmark dataset

We also benchmarked NeuroFold against various deep learning models using the Spearman rank correlation (Figure 5). The Spearman rank correlation with experimentally validated activity/fitness was benchmarked against each model’s output on an input sequence. NeuroFold showed a 0.4 Spearman rank correlation with activity/fitness (Table 1). The next best model was ESM-MSA with a 0.33 Spearman rank correlation, which is a 17.5% decrease compared to NeuroFold. Interestingly, some models exhibited very low correlations (e.g. ESM-1v and AF2), further reflecting the strength of the NeuroFold architecture in robustly predicting protein fitness as determined by the collected experimental dataset. The Spearman correlation is another useful metric to use in conjunction with accuracy, and the aforementioned metrics as a broad criterion for evaluating a model’s consistent ability to discriminate against unfit proteins (Notin et al., 2022). For the current study, however, we prioritized the optimization of precision and f1-score at the cost of correlation, as optimizing precision leads to more immediate practical applications. Overall, NeuroFold’s performance appears to be superior on ProteinGym datasets compared to TAPE datasets. Performance is best on TPMT_HUMAN_Matreyek_2018 (Matreyek., 2018) and worst on TAPE_HEEH_rd3_0872 (Rocklin et al., 2017). It is unclear whether the low performance on certain datasets is due to experimental error, data curation error, or an inability of NeuroFold to generalize to those datasets.

**Figure 5.**
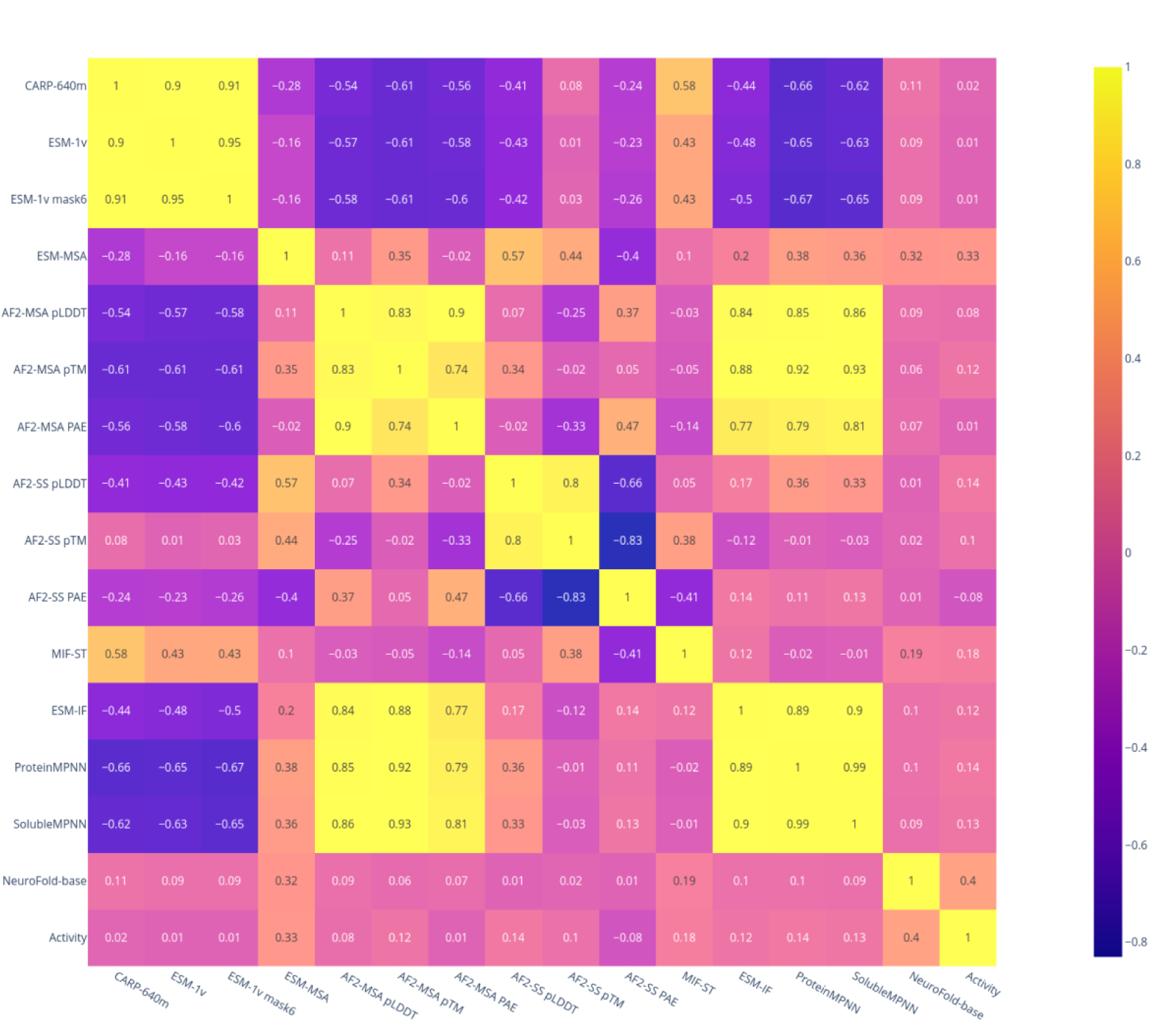
Spearman rank correlation matrix of diverse protein deep learning models. Different models (including NeuroFold) were tested on a collated dataset of 116,321 experimentally validated proteins from 38 protein families/taxa. Different outputs from several different models and algorithms were evaluated for their predictive power of protein fitness based on experimentally confirmed metrics. This correlation matrix highlights the Spearman rank correlation between all results and fitness. ESM-1v mask6 is ESM-1v (Meier et al., 2021) with every 6th amino acid masked, which has been demonstrated to improve discrimination ability (Johnson et al., 2023). SolubleMPNN is ProteinMPNN trained on a dataset of soluble proteins only.

### Experimental Validation

#### Generation of β-Lactamase Variants Using NeuroFold

As a proof-of-principle, we sought to test NeuroFold’s performance in generating variants of a well-characterized and clinically important enzyme. β-lactamase from *Escherichia coli* is a small, monomeric protein with an ⍺-β fold, which catalyzes the hydrolysis of the β-lactam ring of β-lactam antibiotics such as penicillins and cephalosporins (Herzberg and Moult, 1987; Strynadka et al., 1992). This simplicity, in addition to the lack of cofactor required for structure and/or catalysis, makes β-lactamase an ideal candidate for variant production and testing. The active site of β-lactamase is composed of a set of conserved residues implicated in substrate binding and catalysis, including Ser70, Lys73, Ser130, Asn132, Glu166, Asn170, Lys234, Ser235, Gly236 and Ala237 (standard numbering scheme from Ambler et al. (1991) for class A β-lactamases). Ser70 is the nucleophile, directly participating in catalysis and forms the oxyanion hole with Ala237 to stabilize the negative charge formed by the tetrahedral intermediate during acylation and deacylation (Strynadka et al., 1992; Fisher and Mobashery, 2009). Lys73, Asn170, Glu166 and a water molecule are thought to activate Ser70 by abstracting a proton (Herzberg and Moult, 1987; Adachi et al., 1991; Delaire et al., 1991; Escobar et al., 1991; Strynadka et al., 1992; Damblon et al., 1996; Minasov et al., 2002; Meroueh et al., 2005). Ser130 shuttles protons between Lys73 and the leaving group nitrogen (Strynadka et al., 1992). Asn132, Lys234, Ser235 and Gly236 are involved in substrate binding and transition state stabilization (Strynadka et al., 1992; Delmas et al., 2010; Fonseca et al., 2012). Mutations at these residues have all been demonstrated to reduce activity (Palzkill, 2018).

ProteinMPNN was used to generate 4,096 enzyme variants of β-lactamase (PDB ID: 1BTL) using the Neurosnap platform (Neurosnap Inc., 2022; https://neurosnap.ai/). The resultant variants were then filtered using NeuroFold to produce a set of potential candidates. Of these potential candidates, four were randomly selected (see Table S3) for experimentation and their sequence alignment and predicted topology are presented in Figure 6. Three of the variants (i.e. Preserved Variants or PV) had their catalytic sites fixed during the ProteinMPNN sampling process. The catalytic sites were fixed in order to bias ProteinMPNN as well as to maximize the probability of obtaining functional enzymes. This step could be unnecessary as even samples without catalytic site preservation (i.e. Non-Preserved Variant or NPV) subsequently demonstrated partial conservation of the active site residues, with mutations similar to the variation seen across class A β-lactamases.

**Figure 6.**
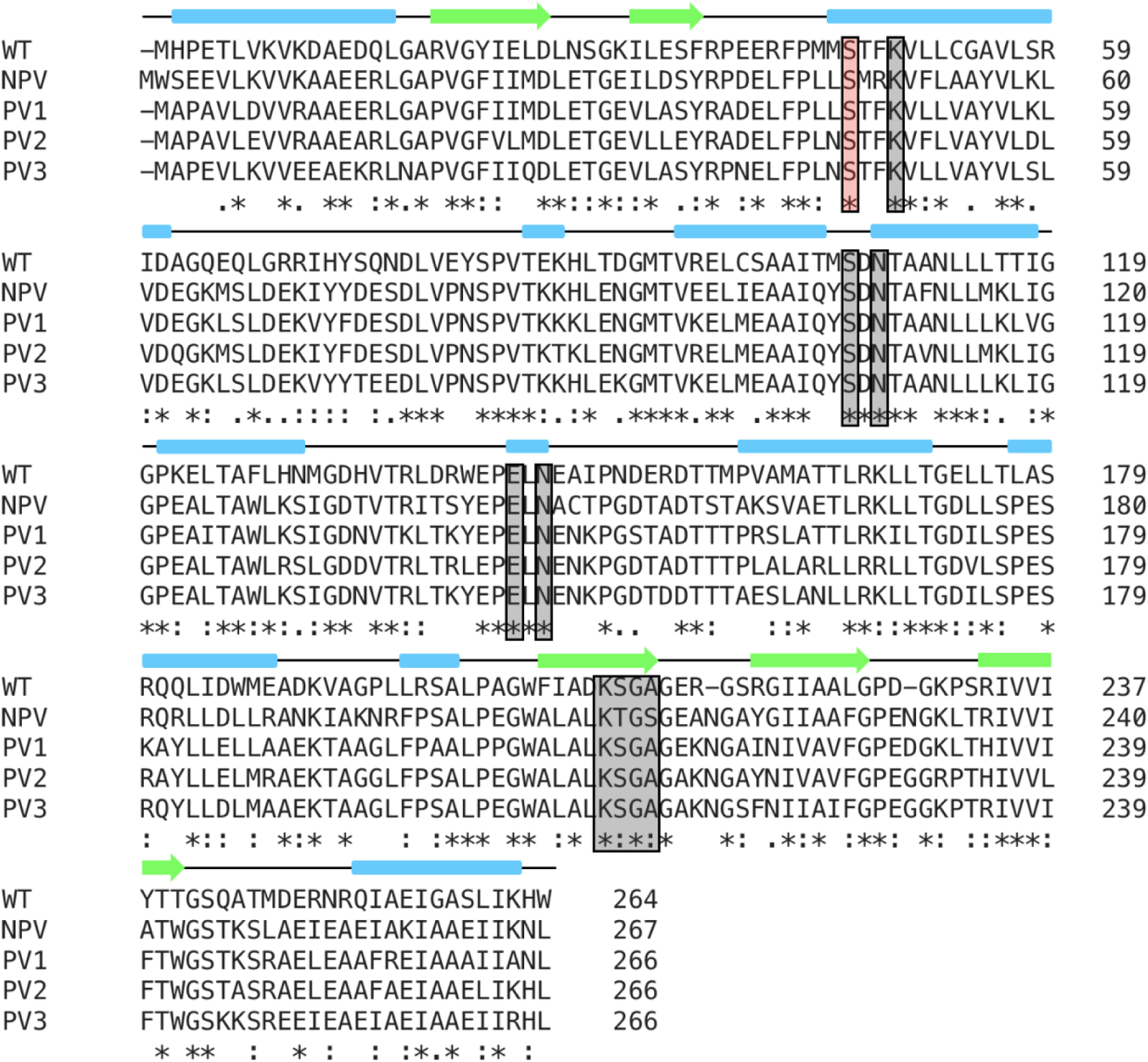
Amino acid sequence alignment of β-lactamase and variants designed by NeuroFold. CLUSTAL multiple sequence alignment by MUSCLE (3.8) using the EMBL-EBI platform (Edgar, 2004). An * (asterisk) indicates positions which have a single, fully conserved residue. A : (colon) indicates conservation between groups of residues with strongly similar properties. A. (period) indicates conservation between groups of residues with weakly similar properties. Secondary structure is mapped based on the crystal structure of WT β-lactamase (PDB: 1BTL) and AlphaFold2 predictions for β-lactamase variants (NPV - Non-Preserved Variant; PV - Preserved Variant). Blue blocks represent ⍺-helices and green arrows represent β-strands. Gray highlights correspond to active site residues that were optionally preserved; the catalytic serine residue is highlighted in red.

Remarkably, all variants selected for experimental validation exhibited sequence identities to the wildtype enzyme ranging from 49% to 56% (Figure 6) and mutations were evenly distributed throughout the entire protein sequence. Despite low sequence identity, the predicted secondary and tertiary structures of the variants were nearly identical (Figure 6 and Figure 7A). The amino acid composition, predicted pI (5.14 - 5.94) and predicted molecular weight (29.3 - 29.9 kDa) were also very similar (Table S4), thus demonstrating that the NeuroFold pipeline is effective at producing variants with low sequence identity, while conserving important physicochemical characteristics.

**Figure 7.**
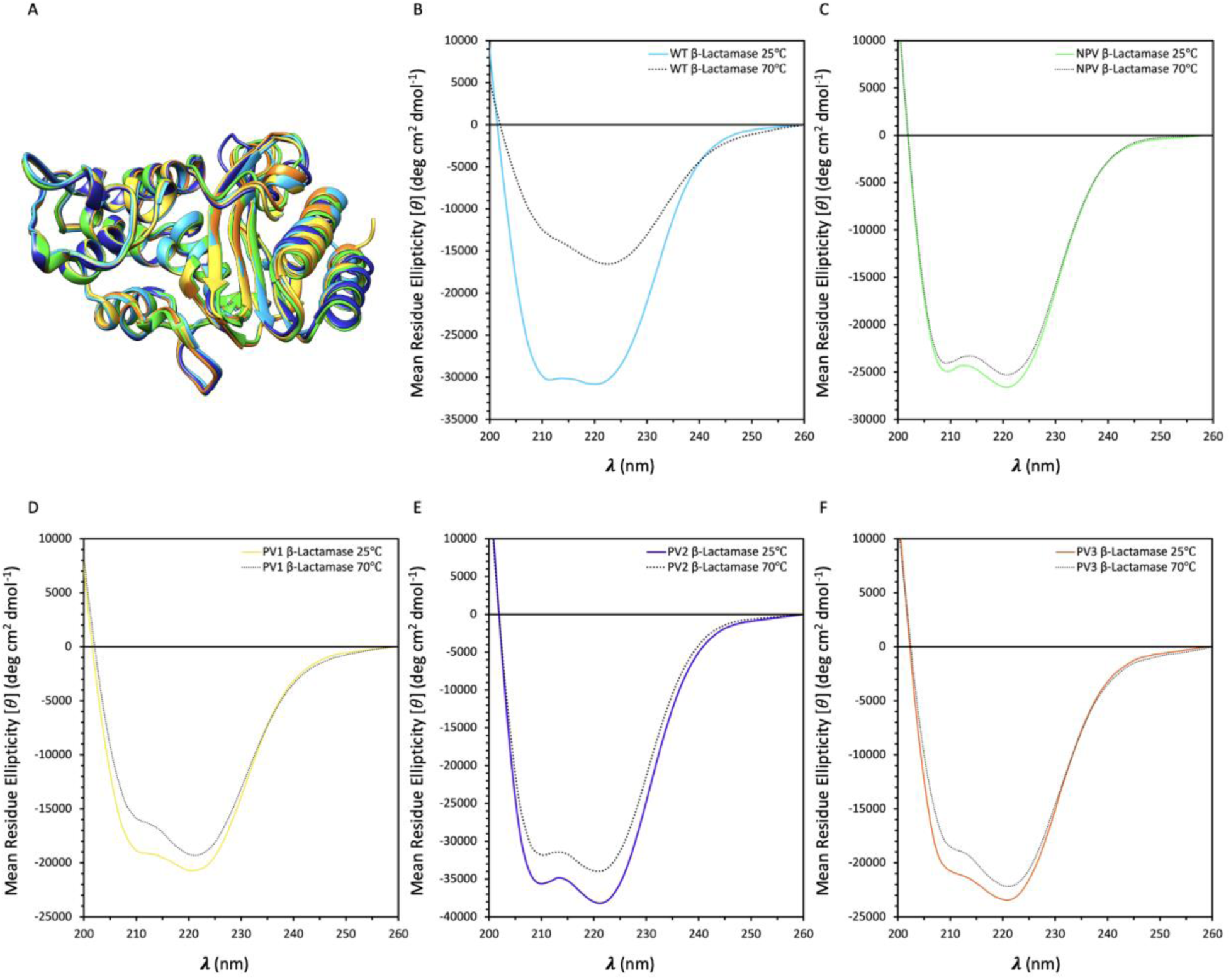
Biophysical Characterization of WT and Variant β-Lactamase Structure and Stability. (A) Structural alignment of WT β-Lactamase (PDB #1BTL) and AlphaFold2 predicted structures for β-Lactamase variants using Chimera MatchMaker, which are in close agreement at the backbone level. (B-F) Far-ultraviolet circular dichroism (CD) spectra at 25℃ (solid and coloured) and 70℃ (dotted and black) for WT (B), non-preserved variant (NPV) (C), preserved variant 1 (PV1) (D), preserved variant 2 (PV2) (E) and preserved variant 3 (PV3) (F) β-Lactamases. All spectra indicate close agreement of secondary structure (⍺-helical), where WT β-Lactamase structure is destabilized at 70℃, marked by a large reduction in the relative ellipticity at 222 nm compared to the variant β-Lactamases.

#### Biophysical Characterization of β-Lactamase Variant Structure and Function

The wild type (WT) β-lactamase and four candidate variants were recombinantly expressed in *E. coli* and purified using immobilized metal affinity chromatography (IMAC) from the soluble fraction. The variants were purified with comparable yields to the WT and migrated similarly when analyzed by SDS-PAGE, with molecular weights of approximately 30 kDa (Figure S2). Each variant had ⍺-helical secondary structure comparable to the WT, as indicated by the far-UV circular dichroism (CD) spectra (Figure 7B-F) with characteristic minima at 208 and 222 nm, apparent maxima below 200 nm, and in agreement with structural predictions using AlphaFold2 (Figure 7A). Light scattering and flattening effects did not influence or distort the spectra within the range of wavelengths shown in Figure 7B-F. Preliminary stability data using far-UV CD spectroscopy above the melting point of WT β-lactamase (70℃) suggests that the variants are more thermostable than the WT, indicated by a reduction in the ellipticity at 222 nm for the WT relative to the variants (Figure 7B-F). This data supports that NeuroFold is capable of generating variants that maintain the structure and increase the stability of the WT protein backbone.

The β-lactamase activity assay was used to assess the enzymatic activity of the variants relative to the wild type (Figure 8A and B). The WT exhibited a specific activity of 6.28 a.u. min^-1^ mg^-1^ at 413 nm for the substrate penicillin G. The non-preserved variant (NPV) exhibited a specific activity of 4.58 a.u. min^-1^ mg^-1^ at 413 nm for the substrate penicillin G. The preserved variants (PV1-3) exhibited specific activities of 0.628, 2.56 and 0.048 a.u. min^-1^ mg^-1^ at 413 nm for the substrate penicillin G, respectively. The NPV, which possessed mutations to some active site residues (A212S and S210T), demonstrated 73% relative specific activity to the WT. PV2 exhibited 41% relative specific activity to the WT, whereas PV1 and PV3 exhibited relative specific activities below 10% compared to the WT. This unpredictability remains one of the major limitations of rational protein design, as it is often the case that mutations far from the active site of an enzyme can have measurable effects on activity (Stiffler et al., 2015). Key catalytic residues in the active site of WT β-lactamase reported in the literature (Palzkill, 2018) were aligned to compare their orientation (Figure 8C-G). The active sites showed very close agreement at the backbone and residue level, even for the NPV. However, minor differences in binding pocket shape, electrostatic surface and substrate binding could be observed between the variants and relative to the WT (Figure 8H-L). These changes suggest that, although the active site residues remain unchanged, substrate binding or transition state stabilization could vary considerably between the variants and relative to the WT, which could explain the significant differences in activity. The results illustrate an important trade-off in the practice of enzyme design – changes to sequence which might confer enhanced stability or other favorable properties often come at the cost of decreased catalytic efficiency. Nonetheless, we show that NeuroFold has the ability to generate enzyme variants that maintain varying levels of activity with less than 56% sequence identity.

**Figure 8.**
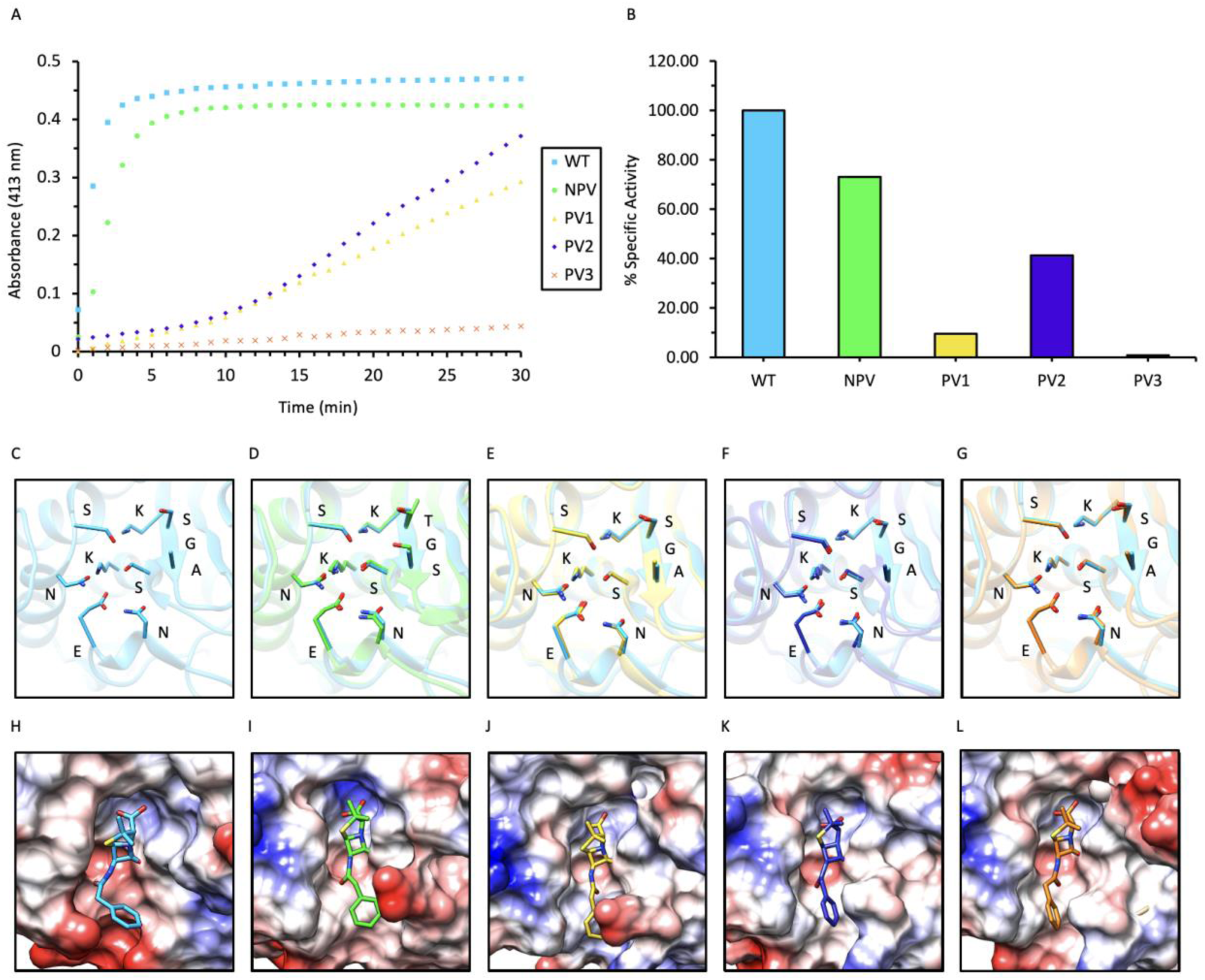
Enzymatic Activity and Computational Characterization of WT and Variant β-Lactamases. (A) Enzyme activity of WT and variant β-lactamases measured as change in absorbance at 413 nm against time (minutes) in the presence of penicillin G substrate and the PAR-2Hg^2+^ complex. An increase in absorbance at 413 nm is a result of the hydrolysis of penicillin G to penicilloic acid, which chelates Hg^2+^ from the PAR-2Hg^2+^ complex, causing a color change from red to yellow. (B) % Specific Activity of variant β-Lactamases relative to the WT determined by linear regression of the tangent line to each curve in (A) at time = 0 min. (C-G) Structural alignment of key active site residues between the WT (C), non-preserved variant (NPV) (D), preserved variant 1 (PV1) (E), preserved variant 2 (PV2) (F) and preserved variant 3 (PV3) (G) β-Lactamases which are in close agreement at the backbone and residue level. (H-L) Binding pocket shape, electrostatic surface and substrate (penicillin G) docking comparisons between the WT (H), NPV (I), PV1 (J), PV2 (K) and PV3 (L) β-Lactamases which indicate minor differences in binding pocket shape, electrostatic surface and substrate binding.

### Concluding Remarks

In this study, we describe NeuroFold, a multimodal model that is able to integrate sequence, structural and evolutionary information to make inferences on the global protein space and to design functional enzyme variants. In particular, we show that NeuroFold can be used to generate divergent versions of the enzyme β-lactamase that exhibit enhanced thermostability and measurable activity levels up to 73% relative to the WT. Despite the remarkable capabilities of NeuroFold, there are certain limitations to consider. Firstly, it should be noted that NeuroFold may encounter challenges when dealing with certain enzyme groups (e.g. orphaned proteins, multimeric proteins, very large proteins) that exhibit incompatible formats not yet supported by the model. Moreover, NeuroFold’s current limitations prevent it from effectively producing variants of large enzymes greater than 2,500 amino acids in length, primarily due to the limitations in VRAM and compute time requirements. Finally, NeuroFold has not yet been sufficiently tested on proteins that contain metal-binding sites, allosteric binding domains, or multimeric enzymes. These limitations highlight areas where further development and improvement of NeuroFold are required to expand its applicability across a broader range of enzymes and molecular configurations.

## Materials and Methods

### β-Lactamase Variant Structure Prediction and Docking with Penicillin G

The structures of each enzyme variant and the wildtype were predicted with AlphaFold2’s ColabFold implementation using the Neurosnap platform (Jumper et al., 2021; Mirdita et al., 2022; Neurosnap Inc., 2022; https://neurosnap.ai/). Molecular docking experiments against penicillin G were conducted using the DiffDock model on the Neurosnap platform (Corso et al., 2023; Neurosnap Inc., 2022; https://neurosnap.ai/). See Table 2 for details.

**Table 2.**
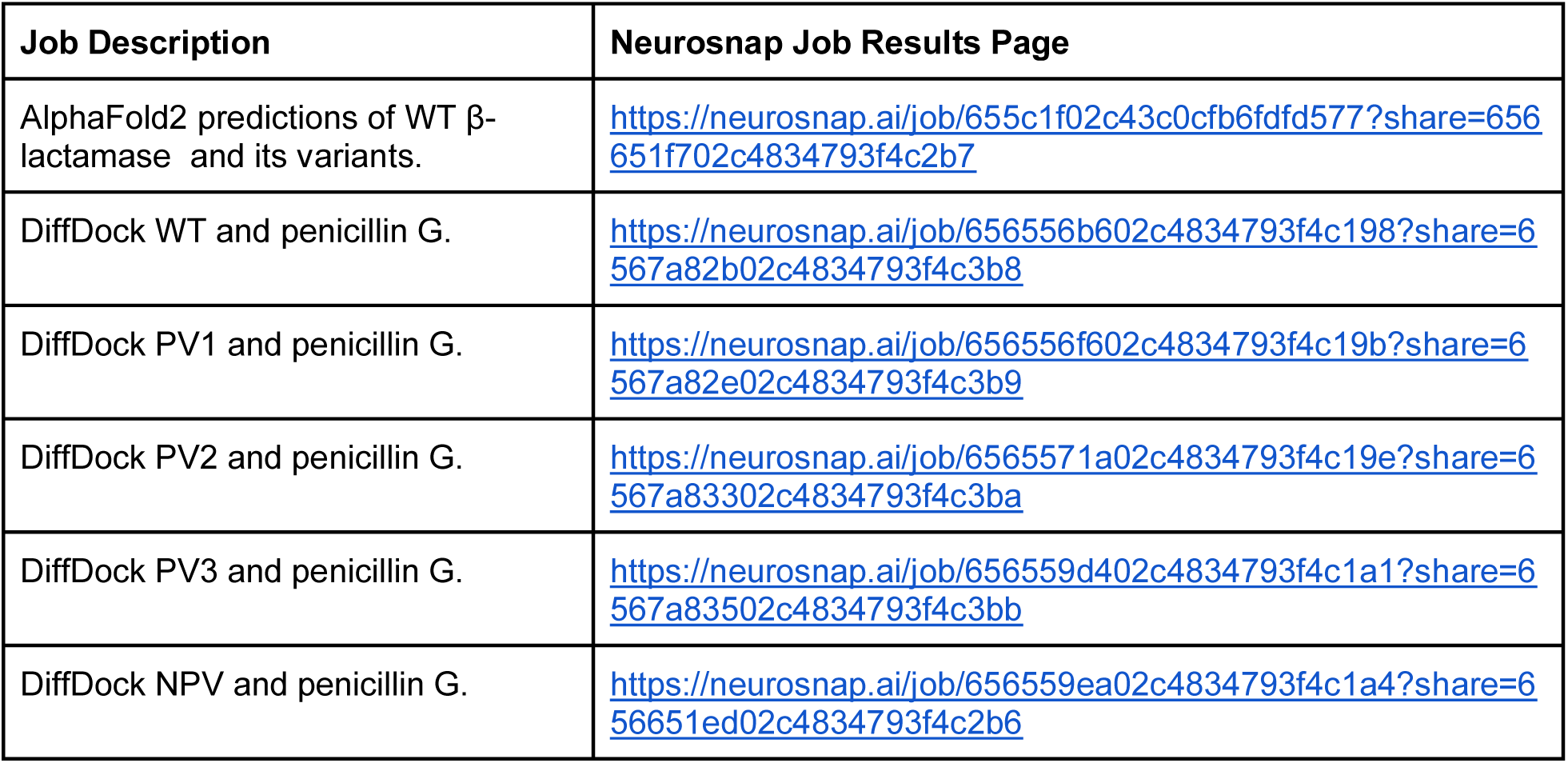
Predicted structures and molecular docking with the target substrate. Structure prediction of the enzyme variants and 1BTL are performed using AlphaFold2 on the Neurosnap platform. DiffDock, a state of the art molecular docking model is used to validate whether the enzyme substrate, penicillin G, interacts with the enzyme’s active site. Each prediction was performed on the Neurosnap platform and thus all the results, inputs, and configurations are available at the job share links.

### Gene Construction

Synthetic cDNA sequences encoding for the wild-type *E. coli* β-lactamase (PDB 1BTL) and 4 variants designed using NeuroFold were synthesized and cloned into pET28(a)+ vectors by Twist Bioscience (https://www.twistbioscience.com/). The endogenous N-terminal transit peptide was removed from the wild-type sequence before variants were generated, so this was not present in any of the constructs. Constructs were designed to include a C-terminal poly-histidine tag fusion for affinity purification and the pET28(a)+ vector provided a kanamycin resistance gene for selection of bacterial clones.

### Protein Expression and Purification

Constructs were introduced into *E. coli* (DH5⍺) and positive clones were isolated, plasmid DNA purified, and plasmids were transformed into *E. coli* BL21 (DE3) LOBSTR cells. Overnight cultures were prepared in LB broth containing Kanamycin from a single colony and incubated at 37°C overnight while shaking at 250 rpm. Overnight cultures were used to inoculate a 1 L culture of LB+Kan and incubated at 37°C for 3-5 hours, until an OD600 of 0.6-0.8 was reached. Expression was induced with 0.1 mM IPTG for 5 hours at 37°C. Cultures were centrifuged for 15 minutes at 4°C and 8,000 x g to pellet the bacterial cells. The supernatant was discarded, and the cells resuspended in lysis buffer (20 mM Tris-HC, pH 8.0, 150 mM NaCl, 5 mM MgCl2, 15 mM imidazole, 1X cOmplete protease inhibitor cocktail (Roche). Cells were incubated in lysis buffer for 1 hour before physical lysis using a Constant Cell Disruptor operating at 20 kPSi. The lysate was centrifuged for 20 minutes at 4°C and 20,000 x g to pellet inclusion bodies and unlysed cells. The supernatant containing the clarified lysate was applied to a pre-equilibrated (with 10 column volumes of lysis buffer) immobilized metal affinity chromatography (IMAC) gravity flow column using Ni-NTA HisPur resin from Bio-Rad with a 1 mL column volume. The clarified lysate was incubated with the resin for 30 minutes at 4°C before collecting the flow-through. The column was washed with 10 column volumes of wash buffer (20 mM Tris-HCl, pH 8.0, 150 mM NaCl, 30 mM imidazole) and the protein was eluted with 10 column volumes of elution buffer (20 mM Tris-HCl, pH 8.0, 150 mM NaCl, 250 mM imidazole) in 1 mL increments. Eluted proteins were desalted to remove imidazole and reduce NaCl to 50 mM (20 mM Tris-HCl, pH 8.0, 50 mM NaCl) using Econo-Pac 10DG Desalting Columns (Bio-Rad), and all fractions were analyzed by Bradford protein assay and SDS-PAGE (4% stacking and 12% resolving gels, stained by Coomassie Brilliant Blue).

### β-Lactamase Activity Assay

A colorimetric assay for β-lactamase activity using penicillin G and the PAR-2Hg^2+^ complex was performed as described by Lee et al. (2017). A solution containing PAR (20 µM), Hg^2+^ (40 µM) and penicillin G (100 µM) in Tris-HCl buffer (20 mM, pH 8.0) was incubated for 40 minutes. After the addition of protein (to a final concentration of 3 µM), the absorbance of the solution was recorded in 1 minute increments for 30 minutes at 413 nm using a BioTek Synergy HT spectrophotometer to determine the change in absorbance/minute per mg of protein.

### Circular Dichroism (CD) Spectroscopy

Far-UV CD spectra of the WT β-lactamase and the 4 variants (3-5 µM) in CD buffer (20 mM Tris-HCl, pH 8.0, 50 mM NaCl) were measured at 25 °C from 190-260 nm using an AVIV 215 spectropolarimeter. Measurements were carried out in a 0.1 cm path-length quartz cuvette at 1 nm resolution. The reported spectra are an average of 9 scans. CD experiments were repeated at 70℃ to monitor structural stability.

## Data & Code Availability Statement

All computed structures, MSAs, and model metrics are freely available for download using the following URL https://neurosnap.ai/neurofold/dataset. Additionally, data can be made available upon reasonable request and by providing the authors with a complimentary lunch.

## Author Contributions

K.A. developed (coded and trained) NeuroFold; conceptualized and performed *in silico* experiments including variant generation, characterization and docking experiments. M.F. conceptualized and performed *in vitro* experiments including wild-type protein selection, construct design, protein expression and purification, activity assays and CD analysis of protein structure and stability. K.A. and M.F. wrote corresponding sections of the manuscript. M.D.S. and C.D.M.C. participated in review and editing of the manuscript in addition to intellectual contributions, funding, and supervision throughout the study. All authors have read and agree to the submitted version of the manuscript.

## Acknowledgements

This research was funded by Natural Sciences and Engineering Research Council of Canada (NSERC); postgraduate doctoral scholarship awarded to M.F. (PGSD3-2021-559542) and Discovery Grants awarded to M.D.S. (RGPIN-2017-05437) and C.D.M.C. (RGPIN-2020-04117) and a Discovery Development Grant awarded to M.D.S. (DDG-2023-00034).

## Ethics Declarations

K.A. is the CEO & Founder of Neurosnap Inc. (https://neurosnap.ai/) and C.D.M.C. sits on the advisory board as a scientific advisor.

## Supplementary Information

**Figure S1.**
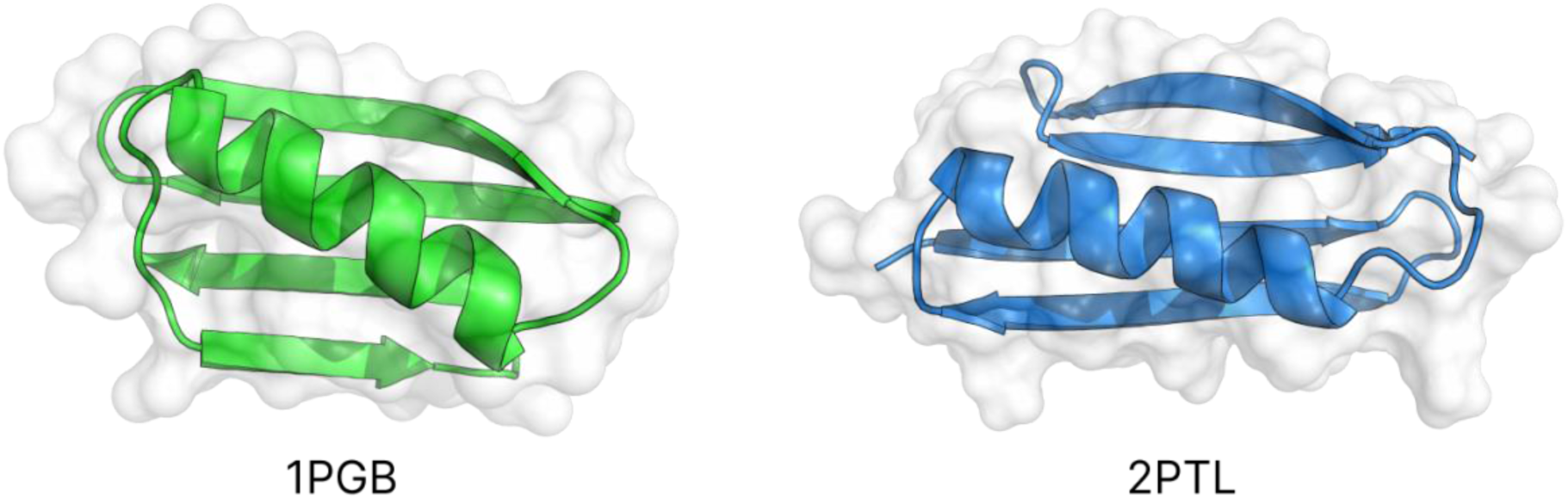
Comparison of 1PGB and 2PTL. The B1 immunoglobulin-binding domain of *streptococcal* protein (1PGB) and the B1 immunoglobulin light chain binding domain of *Peptostreptococcus magnus* (2PTL) have very similar folds despite a pairwise sequence identity of 14% (Koehl & Levitt, 2002).

**Figure S2.**
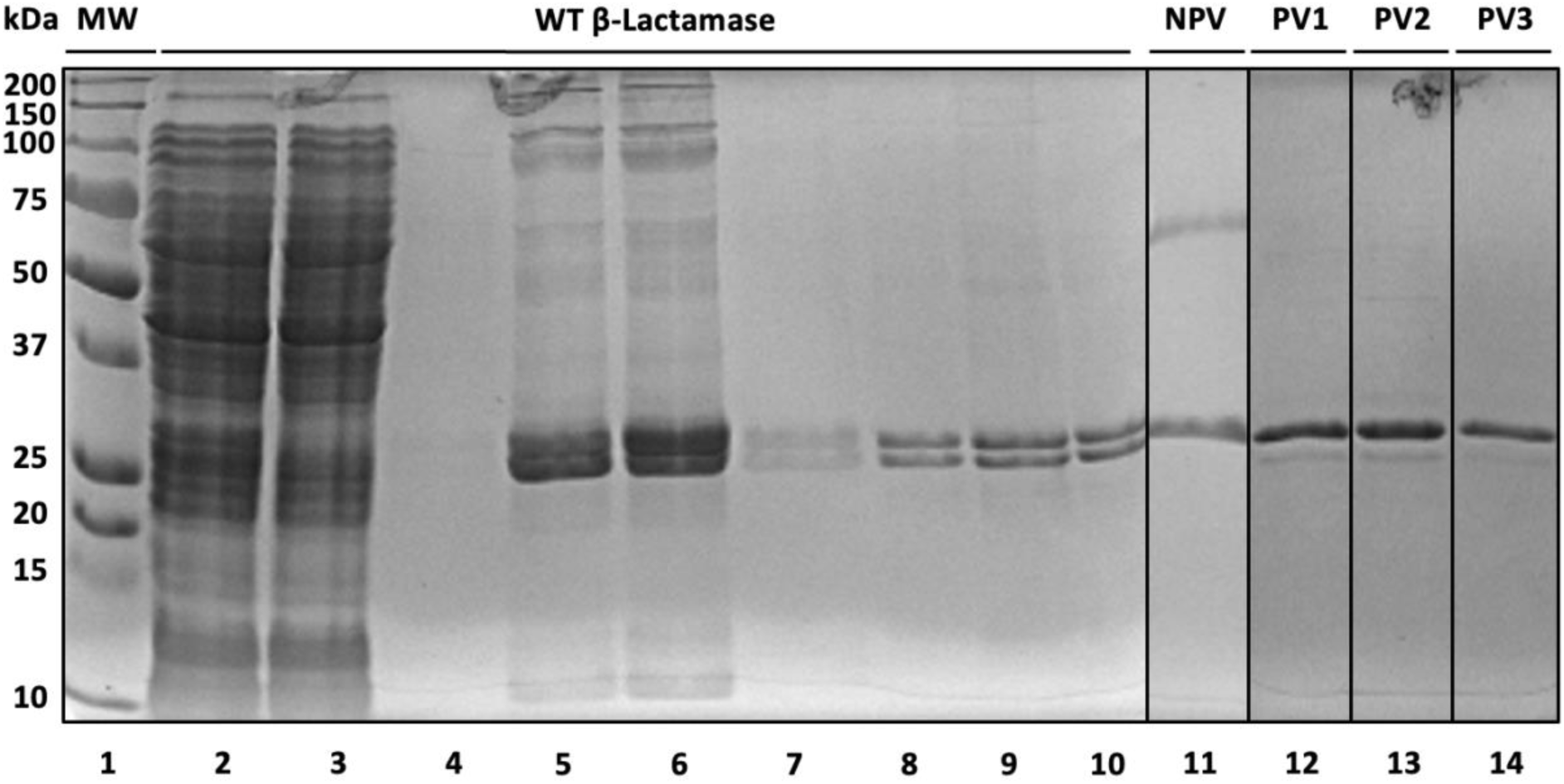
SDS-PAGE Analysis of Expression and Purification of β-Lactamase WT and Generated Variants. (Lane 1) Molecular weight markers in kDa. (Lane 2) *E. coli* whole cell lysate after inducing expression of the WT β-Lactamase. (Lane 3) Flow-Through after applying immobilized metal affinity chromatography (IMAC) with Ni-NTA resin and 5 mM imidazole. (Lane 4) Wash Fraction with 30 mM imidazole. (Lanes 5-7) Elution Fractions with 250 mM imidazole. (Lanes 8-10) Purified WT β-Lactamase after size exclusion chromatography. (Lanes 11-14) Purified β-Lactamase variants non-preserved (NPV), 1, (PV1), 2 (PV2) and 3 (PV3) after size exclusion chromatography. Purified proteins appear at the expected molecular weight (∼30 kDa) between the 25 and 37 kDa markers. Proteins are separated by a 12% resolving gel and visualized by staining with Coomassie Brilliant Blue.

**Table S1.** ProteinMPNN Settings on Neurosnap.

| Setting | Preserved Catalytic Sites | Non-Preserved Catalytic Sites |
| --- | --- | --- |
| PDB | 1BTL | 1BTL |
| Chains | A | A |
| Model Version | original | original |
| Model Type | v_48_020 | v_48_020 |
| Fixed Positions | 70-74, 130-134, 166-171, 234-238 | None |
| # Sequences | 2048 | 2048 |
| Sampling Temperature | 0.01 | 0.01 |

**Table S2.**
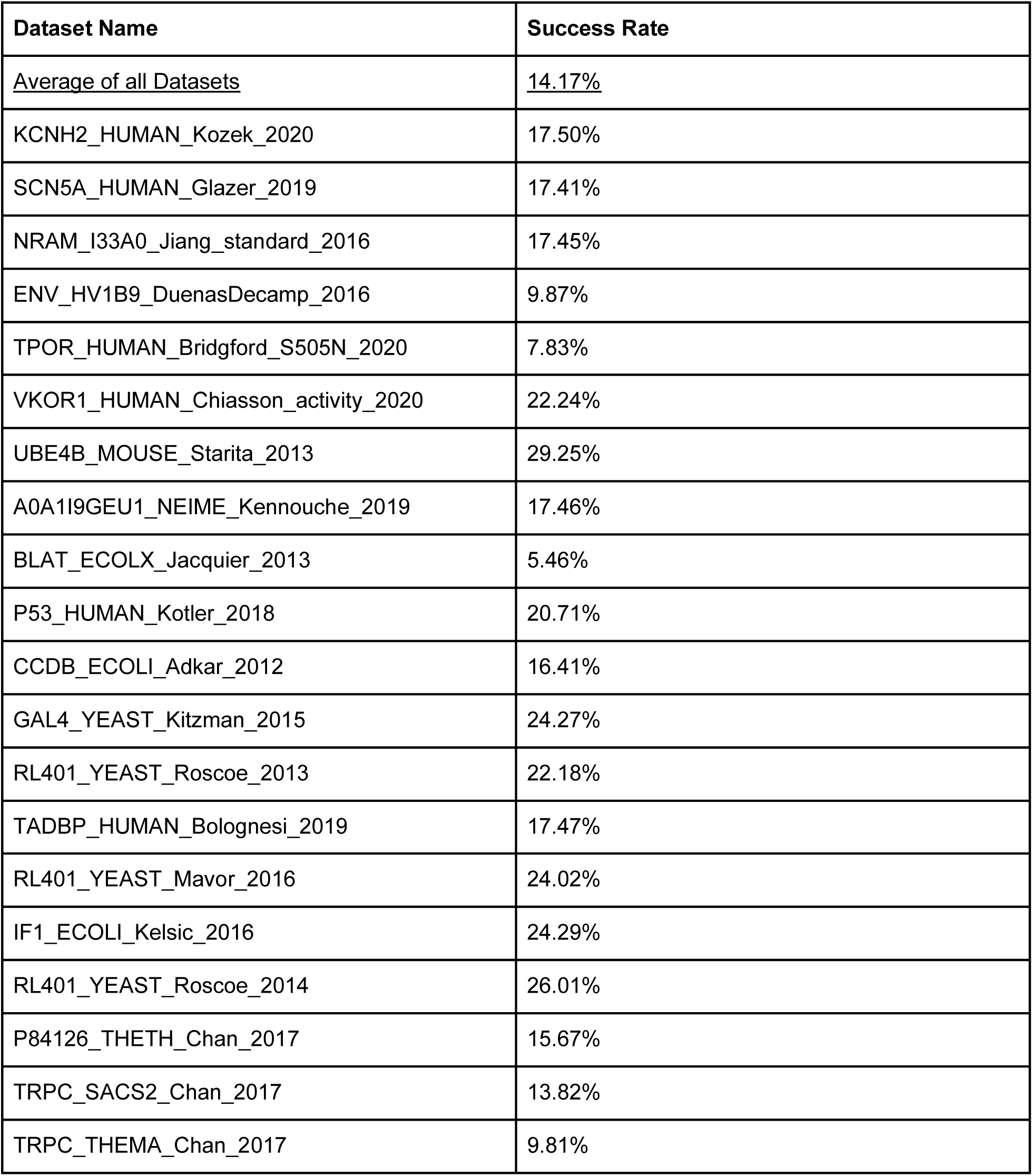
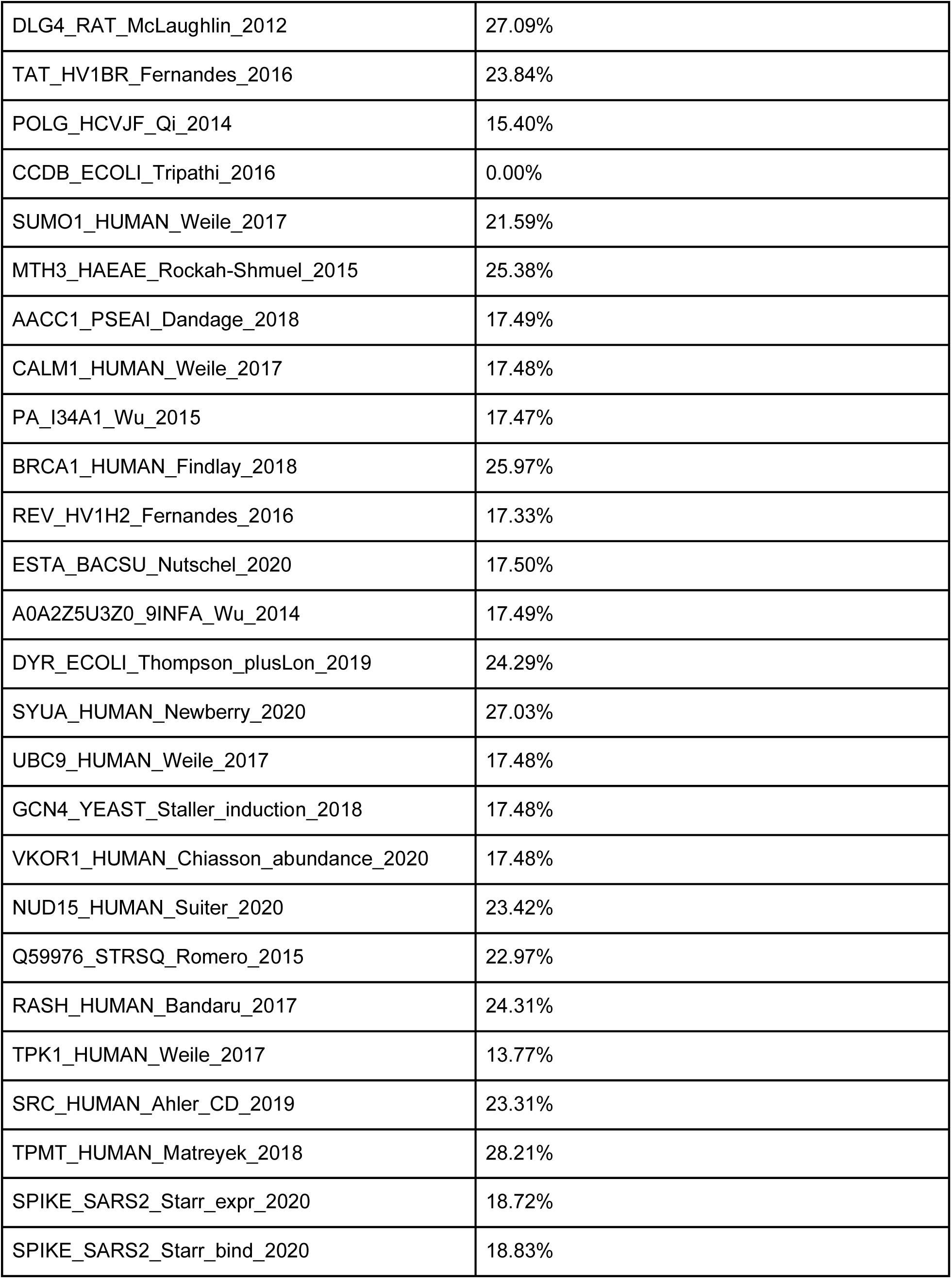
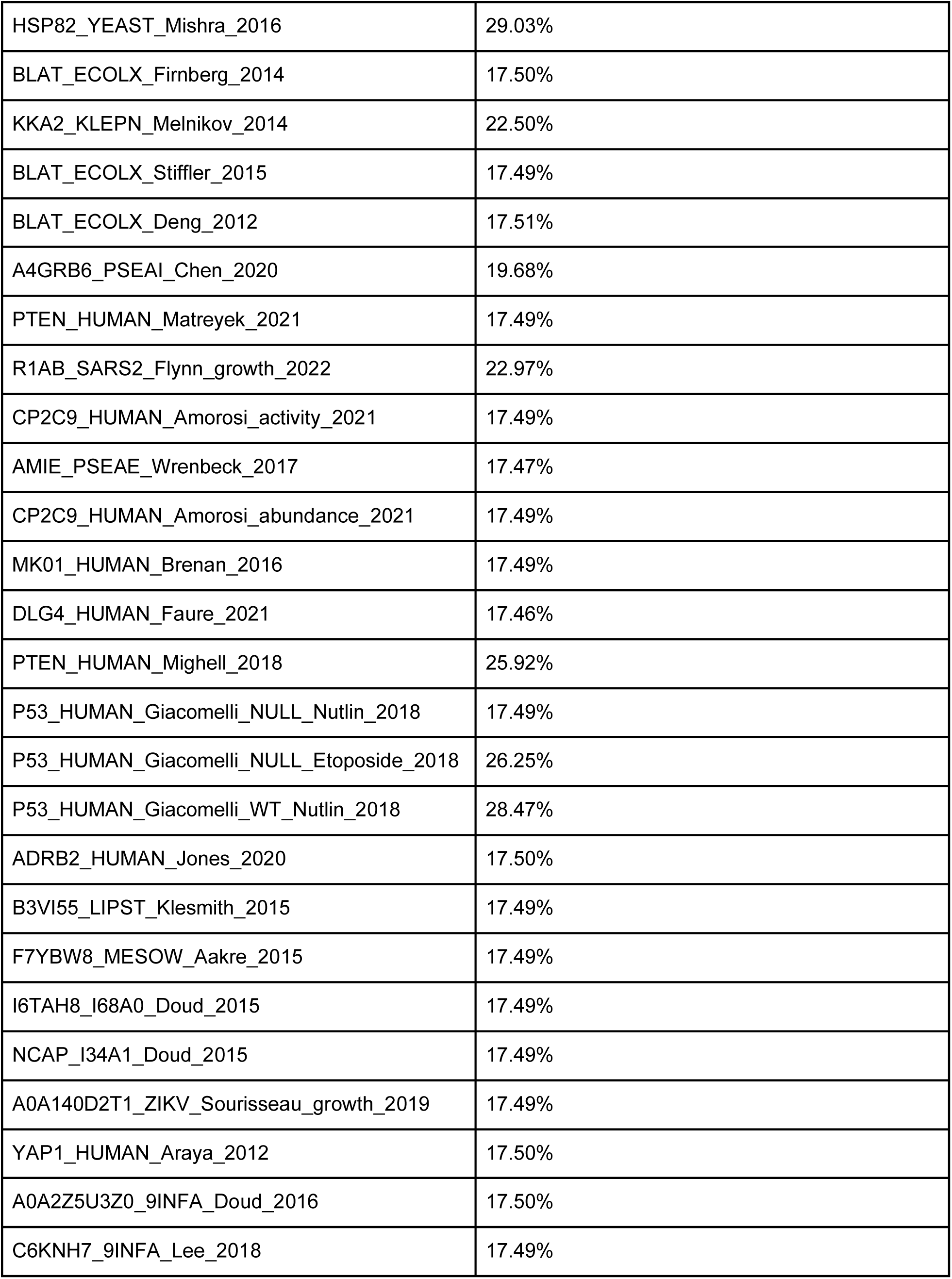
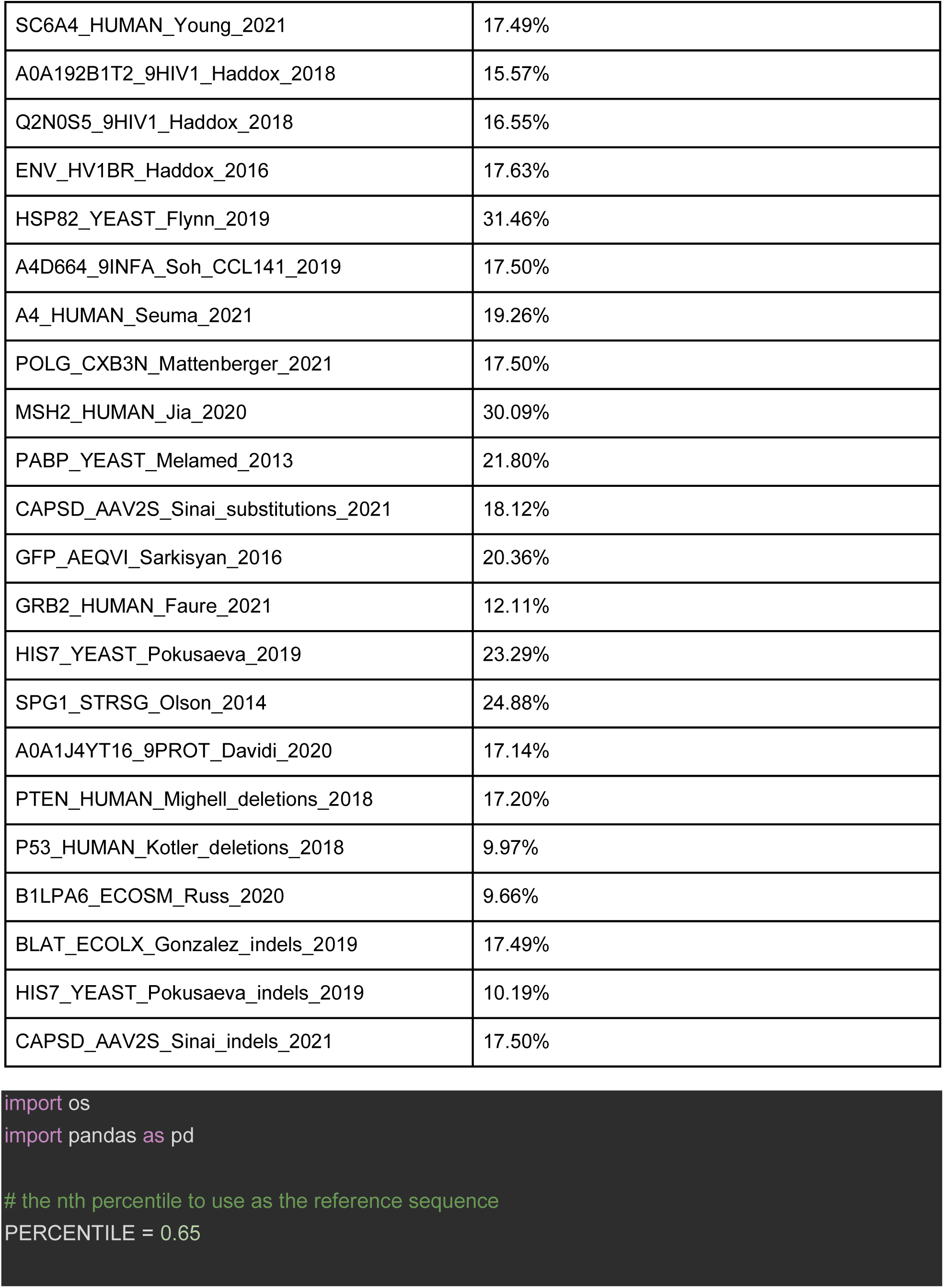
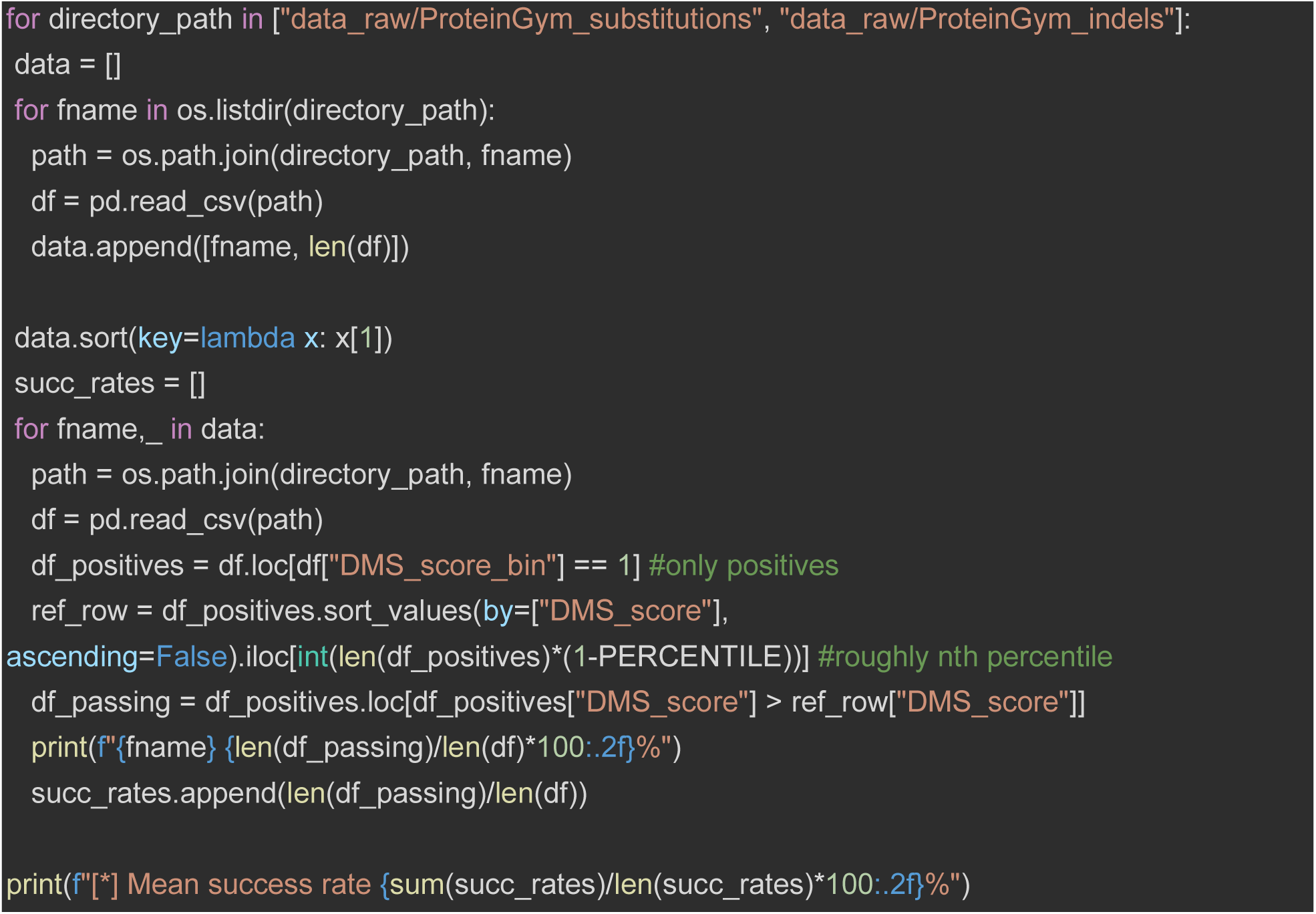
Calculating Average Success Rate of DMS Experiments. We use the ProteinGym dataset (Notin et al., 2022) to calculate the average success rate of a DMS experiment by getting the top 65th percentile of the subset of positives for each dataset to use as the reference sequence. All sequences with a DMS_score greater than the DMS_score of the reference sequence are considered passable. The total number of passing sequences divided by the total number of sequences within that DMS experiment is considered the success rate of that experiment. We calculate the average of the success rates and present them below alongside the average. The python code utilized to produce these numbers is also provided.

**Table S3.**
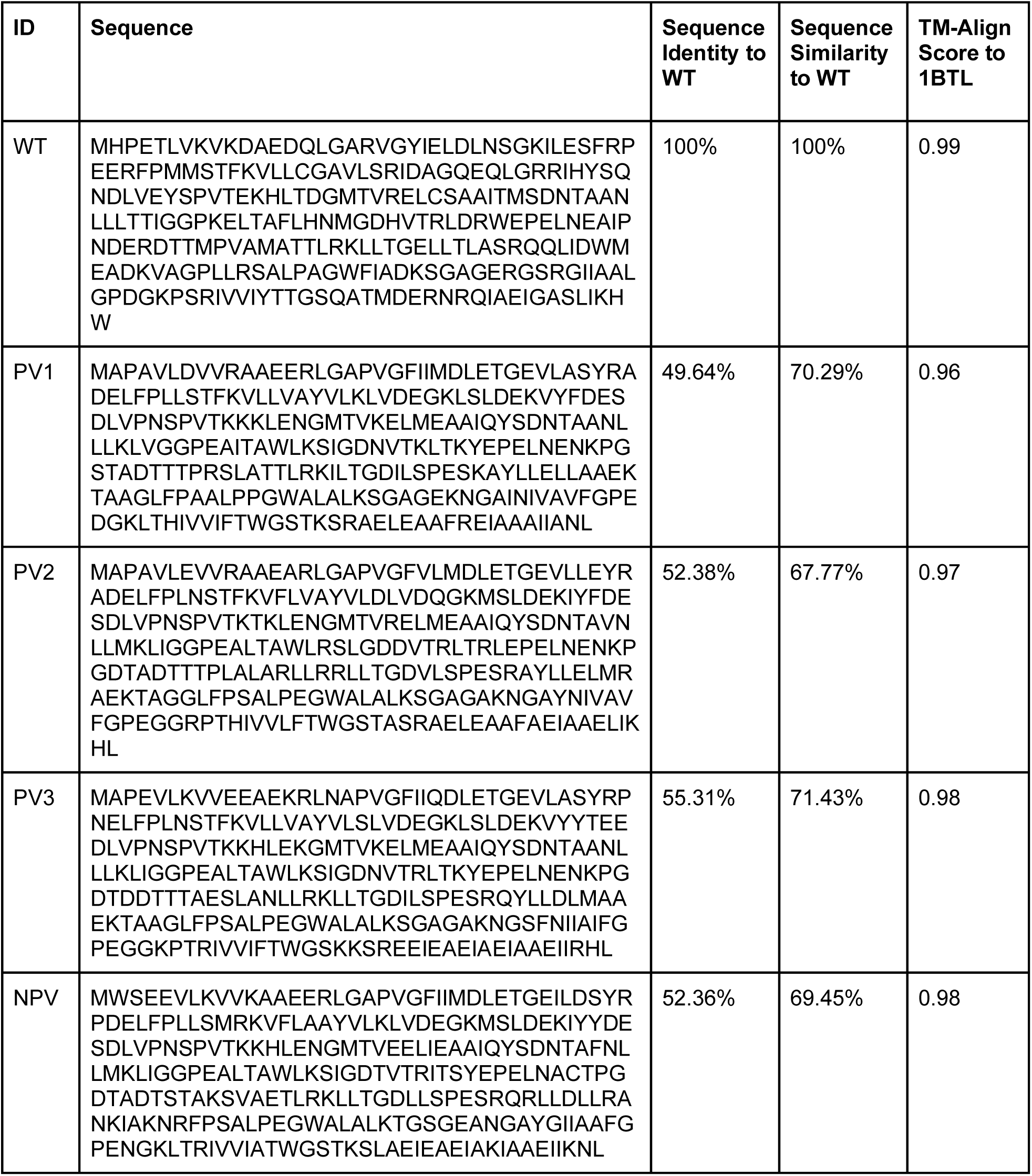
β-Lactamase WT & Generated Variants.

**Table S4.** β-Lactamase WT & Generated Variants. The predicted isoelectric point (pI) and molecular weight (kDa) were determined based on the protein sequence using Expasy ProtParam (Gasteiger et al., 2005).

| ID | Predicted pI | Predicted MW (kDa) |
| --- | --- | --- |
| WT | 5.94 | 29.9 |
| PV1 | 5.44 | 29.3 |
| PV2 | 5.14 | 29.6 |
| PV3 | 5.20 | 29.9 |
| NPV | 5.27 | 29.9 |

**Table S5.** Protein Sequence Identity & Similarity. The settings below were used to produce the sequence identity values within the paper. The online vectorbuilder sequence alignment tool was used with the following settings and URL. **URL:** https://en.vectorbuilder.com/tool/sequence-alignment.html

| Setting Name | Setting Value |
| --- | --- |
| Alignment type | Protein alignment |
| Matrix | EBLOSUM62 |
| Gap penalty | 2.0 |
| Extend penalty | 2.0 |

